# Small RNAs couple embryonic developmental programs to gut microbes

**DOI:** 10.1101/2020.11.13.381830

**Authors:** Hayao Ohno, Zhirong Bao

**Author notes:** Correspondence to (H.O.); (Z.B.).

## Abstract

Maternal exposure to microbes and other environmental factors is known to induce adaptive changes in the progeny, but little is understood about how development of the progeny is changed. We show that *Caenorhabditis elegans* undergoes additional embryonic cell divisions in response to maternal gut microbes such as one producing the biopolymer γ-poly-DL-glutamic acid. The divisions coincide with anatomical changes including left-right asymmetric cell alignment, doubling the association between intestinal cells and primordial germ cells, and improved fecundity. The developmental changes are regulated by soma-to-germline transmission of endogenous RNAi and the miR-35 microRNA family, which targets the LIN-23/CDC-25 pathway. Our findings challenge the widespread assumption that *C. elegans* has an invariant cell lineage that consists of 959 somatic cells and provide insights into how organisms optimize embryogenesis to adapt to environmental changes through epigenetic controls.

---

In many species including human, maternal experience of the environment affects the progeny in terms of metabolism, pathogen and stress resistance, as well as behavior, where environmental cues converge onto DNA/chromatin modifications and small RNAs in the germline to generate and transmit the epigenetic information (Cavalli and Heard, 2019). Embryogenesis is known to respond to maternal environment, such as diapause upon starvation (Renfree and Fenelon, 2017). However, despite centuries of research, it is largely obscure whether, and if so how, the embryo changes its developmental programs in response to environmental factors.

The nematode *C. elegans* is renowned for its invariant cell lineage with exactly 670 cell divisions during embryogenesis (Sulston et al., 1983). To precisely characterize the effects of environmental factors on embryogenesis, we extensively traced the cell lineage in *C. elegans* embryos and analyzed the developmental behaviors of the cells under various environmental conditions using time-lapse 3D imaging (Bao et al., 2006). We found an intriguing phenomenon: when *C. elegans* mothers had experienced harmful microbes such as *Microbacterium nematophilum* CBX102 (Hodgkin et al., 2000) and *Enterococcus faecalis* OG1RF (Darby, 2005), the number of intestinal nuclei was increased in their embryos (Figs. 1A, 1B, and 1C). The extra cell nuclei were generated by cell divisions but not by binucleation of cells because they were surrounded by cell membrane (40/40 nuclei, Fig. S1A).

**Figure 1.**
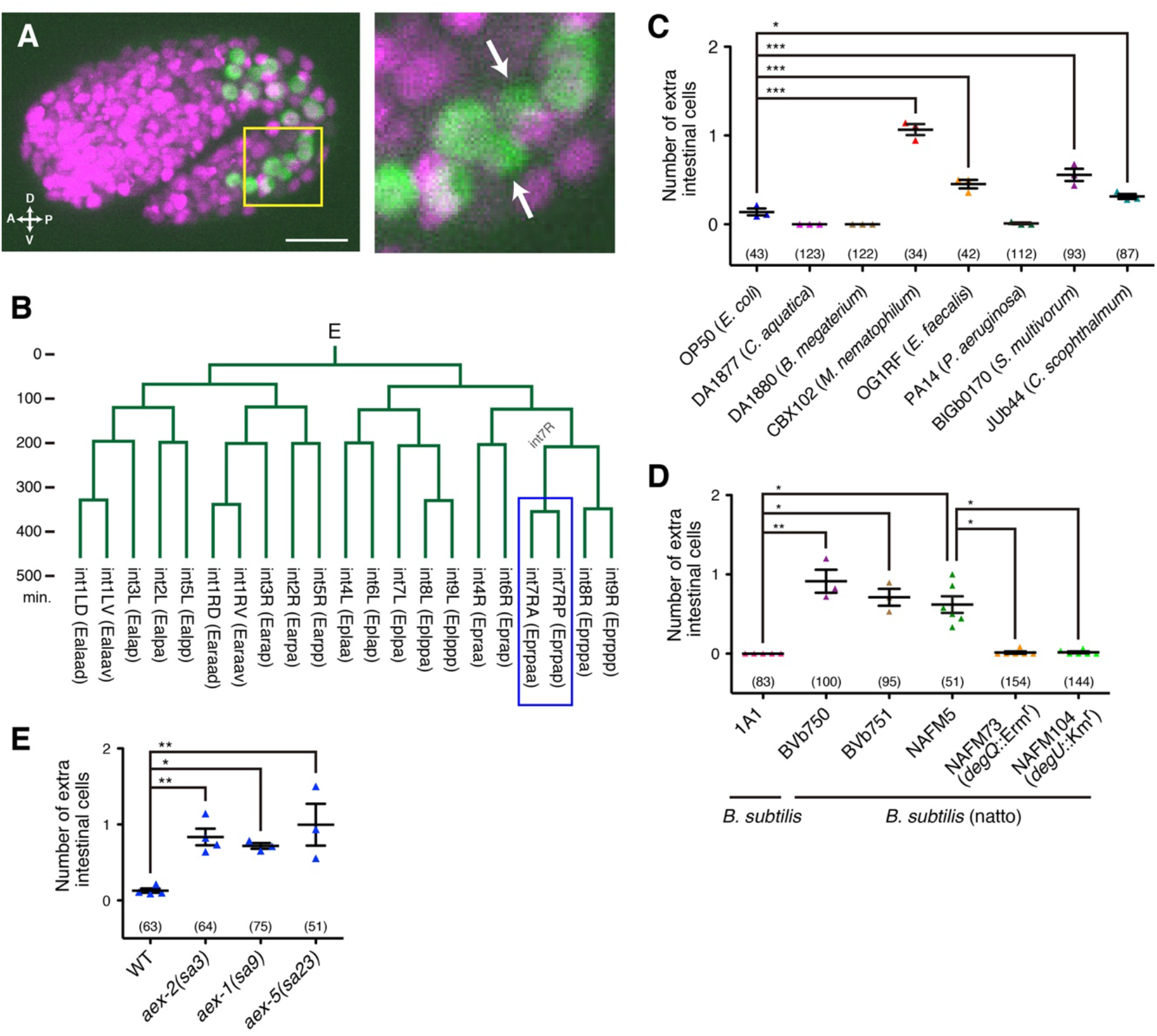
Maternal gut microbes change cell numbers in progeny. (**A**) Representative embryo with one extra intestinal cell, whose mother was exposed to CBX102. Green, intestinal nuclei labeled with *elt-2::GFP;* magenta, all nuclei labeled with mCherry::Histone. A magnification of the boxed region (yellow) is shown in the right panel. White arrows, two nuclei generated by an extra division of int7R. Scale bar, 10 μm. (**B**) The intestinal cell lineage of the embryo shown in (A), acquired at ~one-minute time resolution. The vertical axis is time and a horizontal line indicates a cell division. Blue, extra cell division occurred in int7R (Eprpa). (**C**) Numbers of extra intestinal cells in progeny whose mothers were exposed to various bacterial strains (shown along the X-axis). ***p < 0.001, *p < 0.05 (ANOVA with Dunnett’s post-test). (**D**) Numbers of extra intestinal cells in progeny whose mothers were exposed to *B. subtilis* (1A1), *B. subtilis* (*natto*) (BVb750, BVb751, and NAFM5), or γPGA-nonproducing *B. subtilis* (natto) mutants (NAFM73 and NAFM104). **p < 0.01, *p < 0.05 (Kruskal-Wallis and Dunn’s post-test). (**E**) Numbers of extra intestinal cells of defecation-defective mutants whose mothers were exposed to OP50. **p < 0.01, *p < 0.05 (ANOVA with Dunnett’s post-test). (**C–E**) Bars represent mean ± SEM of at least three independent experiments. Each dot represents the mean from one experiment. The total numbers of progeny scored in all experiments are indicated in parentheses.

The *C. elegans* intestine consists of nine intestinal rings (int1–int9) arranged in an anterior-to-posterior sequence. The first int ring (int1) contains four cells, and the other int rings (int2–int9) each contain two left/right cells (intXL and intXR) (Maduro, 2017). All 20 intestinal cells are generated from the endoderm founder cell named E. Our imaging and subsequent cell tracking in 61 embryos whose mothers had experienced CBX102 showed that in the majority of cases, the extra divisions induced by CBX102 occurred at either int7L (Eplpa) or int7R (Eprpa), or both, increasing the intestinal cells to 21 or 22 (Figs. 1A, 1B, and S1B). A small fraction of the extra divisions (8/66) occurred at int3R (Earap) (Fig. S1B), increasing the intestinal cells up to 23. These extra cell divisions happened almost simultaneously with, or slightly after, the increase in the intestinal cell number from 16 to 20 (Figs. 1B and S1C). Although the axial directions of the extra cell divisions were variable, there is a left-right asymmetric tendency for int7R to divide along the anterior-posterior axis more often than int7L (Fig. S1D).

Lineage changes caused by CBX102 appeared to be specific to the E lineage. We traced the entire cell lineage in three embryos with CBX102-induced extra division in the E lineage up to the 500- to 550-cell stage. Other than the extra divisions in the E lineage, all cell divisions as well as programmed cell deaths up to that stage were normal compared to Sulston’s reference lineage (Sulston et al., 1983) (Figs. S2A–S2C).

Extra intestinal cells were also induced by *Sphingobacterium multivorum* BIGb0170 and *Chryseobacterium scophthalmum* JUb44 (Fig. 1C), both of which are part of *C. elegans*’ natural microbiota that colonize the gut and retard the growth of *C. elegans* (Dirksen et al., 2020), implying that the developmental change also occurs in the *C. elegans*’ natural habitat. *Escherichia coli* OP50, the standard food for *C. elegans* in lab culture but has been reported to be weakly toxic (Darby, 2005; Garsin et al., 2003), had a slight but detectable activity to induce extra intestinal cells (Fig. 1C). Sulston *et al*. noticed that an extra cell was “occasionally” produced in the intestine when they described the entire embryonic cell lineage (Sulston et al., 1983). Our results suggest that their observation is an environmentally induced phenomenon by OP50 rather than intrinsic variability due to less robust control of the lineage. In contrast, no extra intestinal cell was observed on the benign bacteria *Comamonas aquatica* DA1877 (Avery and Shtonda, 2003) and *Bacillus subtilis* 1A1 (Garsin et al., 2003) (Figs. 1C and 1D).

How do the bacteria induce the developmental changes? Given the diversity of the bacterial species, we reasoned that it is more likely to be physiological states of the worm than common chemicals from the bacteria. Highly pathogenic *Pseudomonas aeruginosa* PA14 (Darby, 2005) and the hard-to-eat bacterium *Bacillus megaterium* DA1880 (Avery and Shtonda, 2003) did not induce extra intestinal cells (Fig. 1C), suggesting that neither general health nor nutrient deficiency is a critical determinant. We noticed that all of the bacteria that induced extra intestinal cells are described as viscous (Avery and You, 2012; Powell and Ausubel, 2008) and/or cause “constipation,” “distention,” or “lumen bloating” in the *C. elegans* intestine (Garsin et al., 2001; Hodgkin et al., 2000; Singh and Aballay, 2019; Yuen and Ausubel, 2018) (Fig. S3A). It has recently been suggested that bloating of the intestinal lumen is a “danger signal” to signify microbial colonization (Singh and Aballay, 2019). We examined defecation-defective mutants that are known to cause bloating, namely *aex-1, aex-2*, and *aex-5* (Doi and Iwasaki, 2002; Mahoney et al., 2008), and found that their intestinal cell numbers increased on OP50 (Fig. 1E). Furthermore, restricting the entry of microbes into the gut tube (pharynx, intestine, and rectum) in mothers, but not in fathers, suppressed the increase in intestinal cells induced by CBX102 (Figs. S3B–S3D). We next exposed parental worms to *B. subtilis* (natto). *B. subtilis* (natto) is classified as the same species as the model organism *B. subtilis* but is characterized by the production of a viscous biofilm composed of γ-poly-DL-glutamic acid (γPGA), a major constituent of the sticky strings of the Japanese popular food Natto (Do et al., 2011). *B. subtilis* (natto) strains strongly induced extra intestinal cells in *C. elegans* embryos (Fig. 1D). *B. subtilis* (natto) mutants defective for the γPGA synthesis (Do et al., 2011) did not cause extra intestinal cells (Fig. 1D), indicating that γPGA is required for the induction. *B. subtilis* (natto) also slowed down the growth of worms in a γPGA-dependent manner (Fig. S3E). Taken together, these results raise the possibility that the maldigestion of maternal gut microbes that produce a viscous biofilm and/or are highly proliferative, which is likely to pose a great survival risk to worms (Garsin et al., 2001; Hodgkin et al., 2000; Singh and Aballay, 2019; Yuen and Ausubel, 2018), is an important factor for this developmental plasticity. However, we cannot exclude a potential involvement of a different chemical or physical action(s) of the microbes.

To better understand the effect of maternal gut microbes, we further examined embryonic development and fitness of the progeny whose mothers had experienced CBX102. First, we found that extra intestinal cells are accompanied by changes in the anatomy of the intestine. In embryos that had undergone the extra cell divisions, the right side intestinal cells near the mid-intestine moved anteriorly, resulting in a left-right asymmetric cell arrangement (Figs. 2A, 2B, and S4A); for example, int5R was paired with int4L, instead of the normal int5L (Figs. 2B and 2C) and int5R was even slightly anterior to int4L in ~60% of embryos (Fig. S4B). This left-right asymmetric cell pairing was resolved by the inclusion of int3R in the int2 ring (Fig. S4C). The left-right asymmetry in the directions of the extra int7 division axis (Fig. S1D) seems to be correlated with the left-right asymmetric cell arrangement.

**Figure 2.**
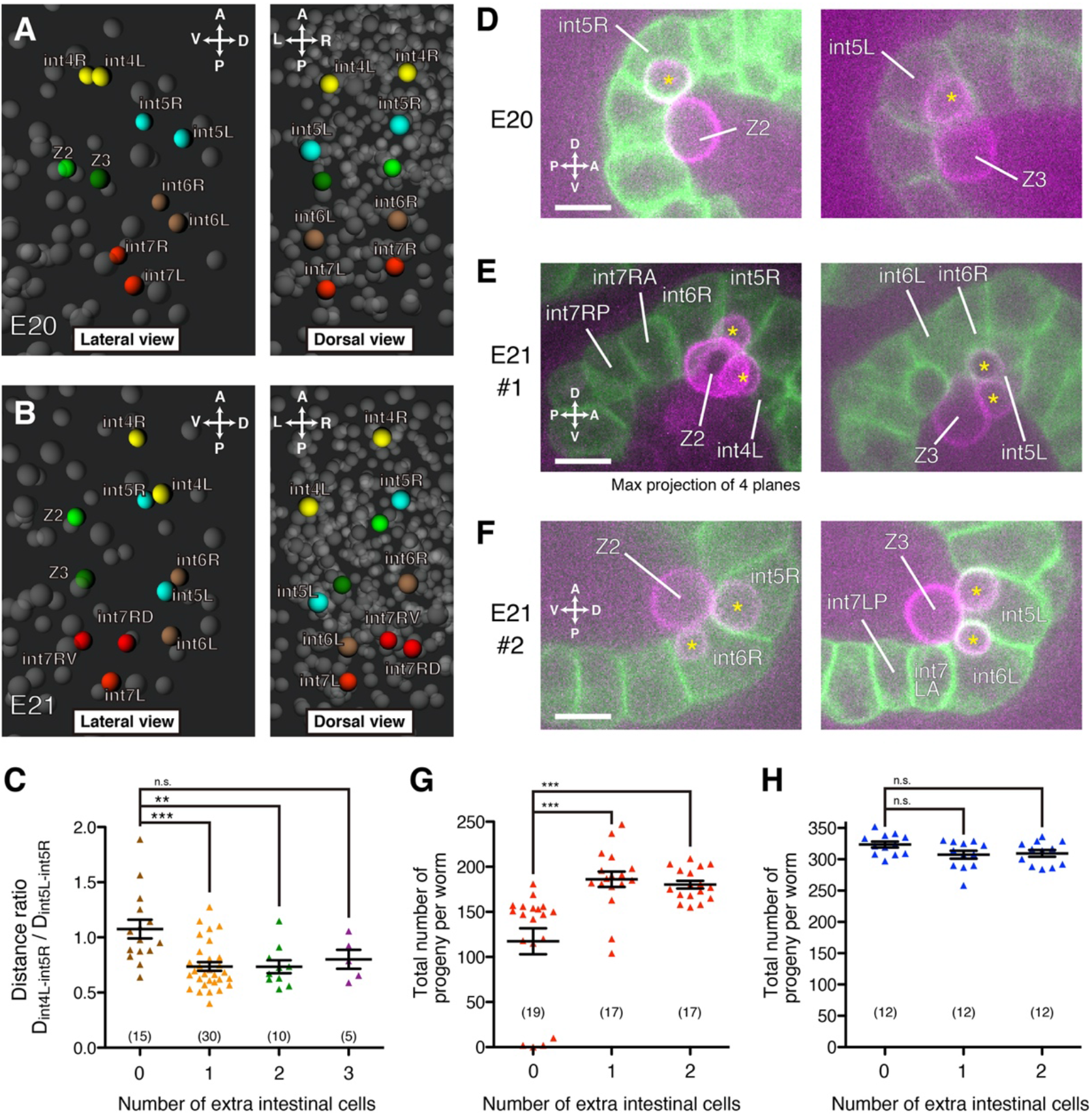
Extra intestinal cells coincide with left-right asymmetric cell pairing, increasing the PGC-intestine association. (**A** and **B**) 3D reconstruction of centers of cell nuclei around the mid-intestine of 1.5-fold stage embryos. An embryo without extra intestinal cell (A, E20) and one with one extra intestinal cell (B, E21), both of which were born to mothers that had been exposed to CBX102, are shown. Left panels, lateral view; right panels, dorsal view. Yellow, int4L/R cells; cyan, int5L/R cells; brown, int6L/R cells; red, int7L/R cells and int7R daughters; yellow-green, Z2; green, Z3. (**C**) Ratios of the distance between int4L and int5R (D_int4L-int5R_) to that between int5L and int5R (D_int4L-int5R_) were measured from 3D images of 1.5-fold stage embryos whose mothers had been exposed to CBX102. ***p < 0.001, **p < 0.01 (ANOVA with Dunnett’s posttest). n.s., not significant. The numbers of embryos are indicated in parentheses. (**D–F**) One representative embryo without extra intestinal cell (D) and two representative embryos with one extra intestinal cell (E and F), whose mothers were exposed to CBX102. Green, intestinal cell membranes labeled with *end-1^prom^::gfp::caax*; magenta, PGC membranes labeled with *mex-5^prom^::mCherry-PH*. Single focal plane images are shown unless otherwise indicated. Left panels, Z2 and its lobes (asterisks); right panels, Z3 and its lobes (asterisks). Scale bars, 10 μm. (**G** and **H**) Numbers of total progeny from worms grown on CBX102 (G) and OP50 (H). Intestinal cell numbers were determined in F1 embryos born to P0 mothers that had been exposed to CBX102, and numbers of F2 progeny of individually cultured F1 worms were counted. ***p < 0.001 (Kruskal-Wallis and Dunn’s post-test). n.s., not significant. Each dot represents one F1 worm. The numbers of F1 worms tested are indicated in parentheses. (**C, G, H**) Bars represent mean ± SEM.

Second, embryos with extra intestinal cells show altered interactions between the intestine and the primordial germ cells (PGCs). From the 88-cell stage to the L1 larval stage, *C. elegans* has two PGCs, namely Z2 and Z3. As in canonical embryogenesis, in embryos without extra intestinal cells, Z2 and Z3 take on an hourglass shape and each insert a lobe into int5R and int5L, respectively (Abdu et al., 2016; Maduro, 2017; Sulston et al., 1983) (Fig. 2D). Through the lobes, PGCs are thought to be nursed by the intestinal cells (Sulston et al., 1983), such as by allowing for clearance of unwanted organelles and other cytoplasmic components (Abdu et al., 2016). In embryos with extra intestinal cells, we found that PGCs altered their morphology and adopted a Mickey-mouse shape by forming an additional lobe with other intestinal cells (Figs. 2E and 2F, 29/30 embryos). The identity of the partner intestinal cells appears to be correlated to the degree of left-right asymmetry of the intestine (Fig. S5).

Third, worms that underwent extra intestinal cell divisions during their embryogenesis show improved fecundity on CBX102 than worms without extra intestinal cells. Worms with extra intestinal cells produced >50% more progeny (Figs. 2G and S6A). In contrast, on OP50 they did not produce more progeny (Figs. 2H and S6B). Their developmental speed on CBX102 was comparable to worms without extra intestinal cells (Fig. S6C), implying that the reproductive fitness is not due to better general health. It is currently unknown if the altered PGC-intestine interaction contributes to the improved fecundity. Nonetheless, the three observations suggest that maternal gut microbes lead to alternative developmental programs with improved fitness.

The *C. elegans* embryo is isolated from the external environment by the eggshell and thus information on microbes is highly likely to be transmitted through the maternal germline. To understand how the information reaches the maternal germline, we investigated the developmental phenotype of mutants of genes implicated in stress responses and found that endogenous RNA interference (endo-RNAi), RNAi caused by organisms’ endogenous small interfering RNAs (Ambros et al., 2003; Ruby et al., 2006), is involved in this plasticity; mutants of *rde-4, eri-1*, and *rrf-3*, all of which are components of the enhanced RNAi (ERI) complex required for endo-RNAi biogenesis (Billi et al., 2014; Grishok, 2013), showed increases in intestinal cells on OP50 (Fig. 3A). This result may be consistent with an increased intestinal cell number caused by RNAi of *eri-5* (Roy et al., 2014), which is another ERI component (Billi et al., 2014; Grishok, 2013). In *C. elegans*, intercellular transport of double-stranded RNAs (dsRNAs) leads to the propagation of RNAi from cell to cell (systemic RNAi), including transmission from soma to germline (Kadekar and Roy, 2019; Kaletsky et al., 2020; Posner et al., 2019). Systemic RNAi requires the dsRNA importer SID-1 (Jose et al., 2009; Shih and Hunter, 2011), the worm ortholog of mammalian SIDT1 and SIDT2, and it has been suggested that the SID-1-dependent intercellular propagation also occurs in endo-RNAi (Kadekar and Roy, 2019; Posner et al., 2019). Mutants of *sid-1* also showed increases in intestinal cells (Fig. 3A). These results suggest that systemic endo-RNAi represses the alternative developmental program.

**Figure 3.**
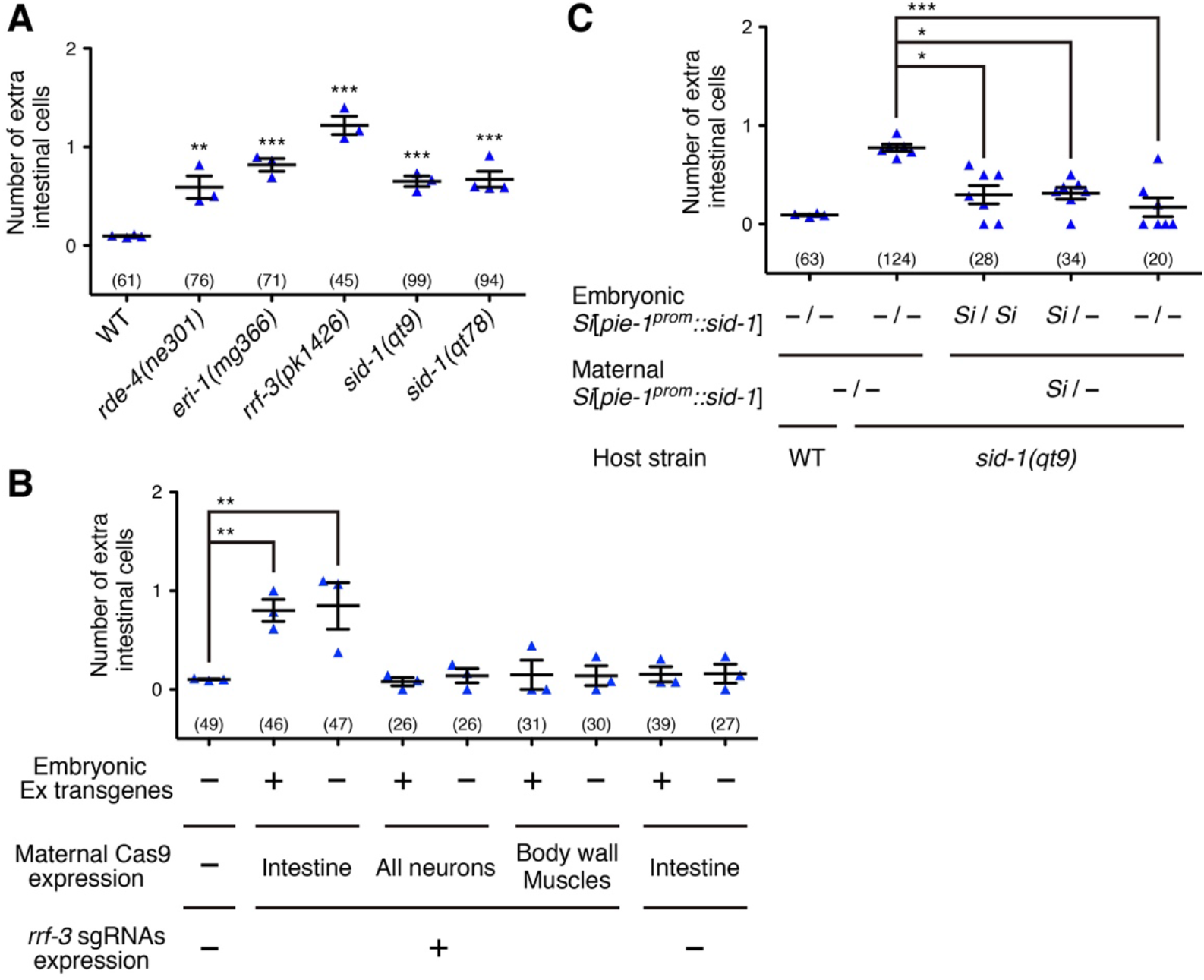
Soma-to-germline transmission of endo-RNAi negatively regulates intestinal cell numbers. (**A**) Numbers of extra intestinal cells of RNAi mutants whose mothers were exposed to OP50. ***p < 0.001, **p < 0.01 (ANOVA with Dunnett’s post-test). (**B**) Cas9 was maternally expressed in a tissue-specific manner with (“+”) or without (“−”) *rrf-3* sgRNAs from extrachromosomal transgenic arrays (Ex) and numbers of extra intestinal cells were counted in array-positive (“+”) and array-negative (“−”, stochastic array loss) embryos. All mothers were exposed to OP50. **p < 0.01 (ANOVA with Dunnett’s post-test). (**C**) The *sid-1* cDNA was expressed specifically in the germline from a single-copy-inserted transgene (*Si*[*pie-1^prom^::sid-1*]) and numbers of extra intestinal cells in progeny segregated from worms heterozygous for the transgene (Si/−) or worms lacking the transgene (−/−) were counted. All mothers were exposed to OP50. ***p < 0.001, *p < 0.05 (Kruskal-Wallis and Dunn’s post-test). (**A–C**) Bars represent mean ± SEM of at least three independent experiments. Each dot represents the mean from one experiment. The total numbers of progeny scored in all experiments are indicated in parentheses.

Conditional knockout of *rrf-3* by somatic CRISPR-Cas9 (Shen et al., 2014) in the maternal intestine, but not that in maternal neurons or body wall muscles, induced extra intestinal cells on OP50 (Fig. 3B), making the intestine a strong candidate as the source of endo-RNAi. Furthermore, the *sid-1* mutant phenotype was rescued by germline-specific expression of *sid-1* driven from a single-copy transgene (*Si*[*pie-1^prom^::sid-1*]) (Fig. 3C), suggesting a role for soma-to-germline transmission of dsRNAs. The rescue effect was observed in transgene-free embryos (−/−) born to heterozygous mothers (*Si*/−) (Fig. 3C), indicating that the maternal action of *sid-1* is important. Taken together, these results suggest that endo-RNAi transmitted from the maternal intestine to the maternal germline negatively regulates the developmental plasticity.

We next focused on microRNAs of the miR-35 family (miR-35–42, hereafter referred to as miR-35^fam^) as potential epigenetic regulators, because their expression is highly enriched in the germline (McEwen et al., 2016) and they positively regulate intestinal cell numbers (Liu et al., 2011). In *mir-35–41(nDf50)* and *mir-35–41(gk262)* deletion mutants (Liu et al., 2011; McJunkin and Ambros, 2014), both of which lack a gene cluster consisting of seven out of the eight family members, CBX102-induced extra intestinal cells were completely suppressed (Fig. 4A). Furthermore, introduction of *nEx1187*, a multi-copy expression array of *mir-35* (Alvarez-Saavedra and Horvitz, 2010; McJunkin and Ambros, 2014), induced extra intestinal cells on OP50 (Fig. 4B). This induction of extra intestinal cells was observed even in array-negative embryos born to array-positive mothers (Fig. 4B), indicating that the increased maternal *mir-35* expression is sufficient for the regulation. Finally, RT-qPCR showed that mature miR-35-3p and miR-40-3p, which are miRNAs expressed from the *mir-35–41* cluster, were increased in the germline when worms had experienced CBX102 or *B. subtilis* (natto) BVb750 (Fig. 4C). miR-35-3p and miR-40-3p were not detected in the *mir-35–41(nDf50)* deletion mutants, indicating the specificity of our qPCR (Fig. 4C). Taken together, these results show that expression level of miR-35^fam^ in the maternal germline rises in response to maternal gut microbes to regulate intestinal cell numbers in the embryo.

**Figure 4.**
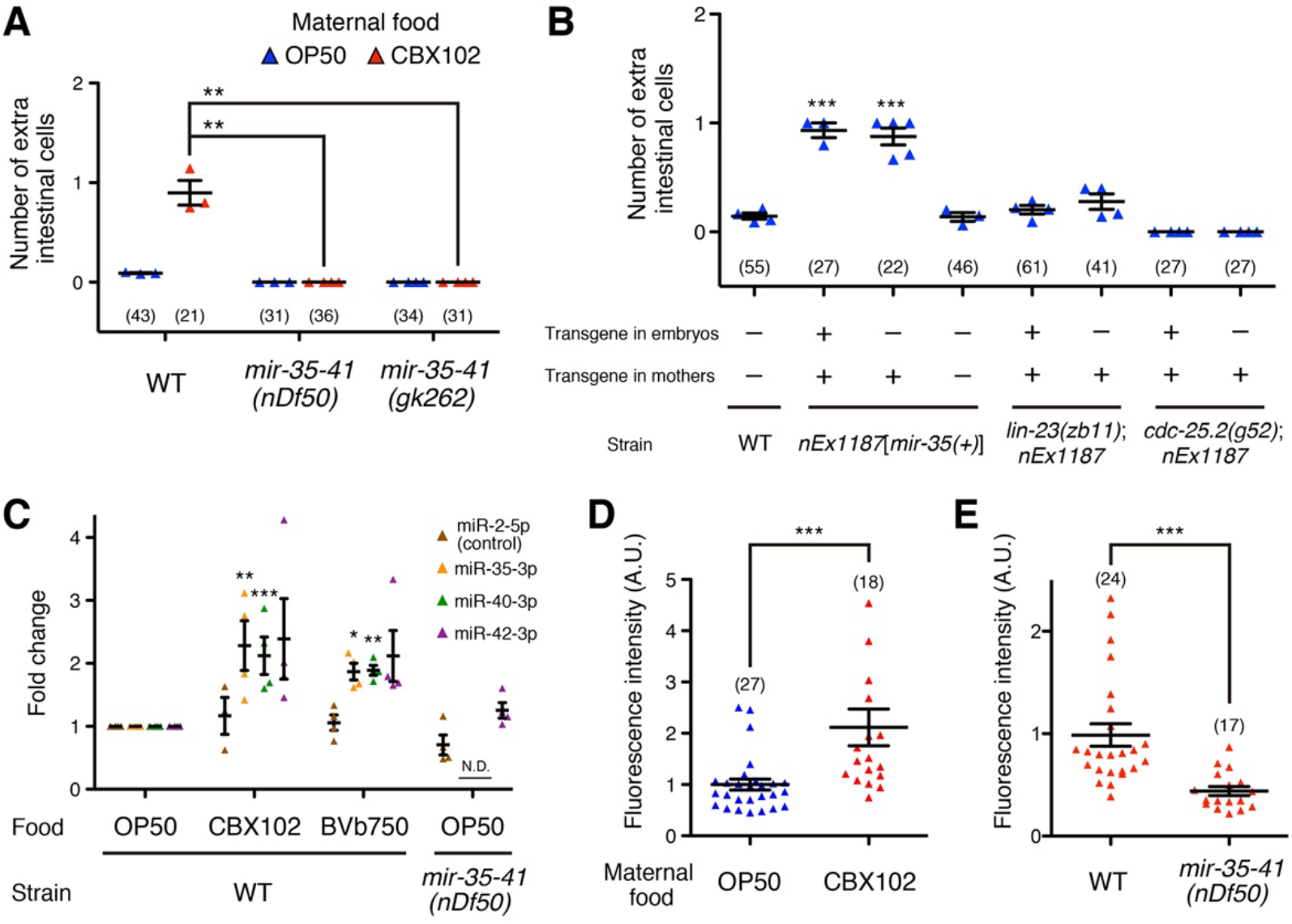
The miR-35 microRNA family is involved in extra intestinal cell divisions. (**A**) Numbers of extra intestinal cells of *mir-35–41* mutants whose mothers were exposed to OP50 or CBX102. **p < 0.01 (Kruskal-Wallis and Dunn’s post-test). (**B**) *mir-35(+)* was expressed from an extrachromosomal transgenic array in the WT, *lin-23(zb11)*, and *cdc-25.2(g52)* backgrounds and numbers of extra intestinal cells were counted in array-positive (“+”) and array-negative (“−” stochastic array loss) embryos. ***p < 0.001 (ANOVA with Dunnett’s post-test, compared with WT). (**A** and **B**) Each dot represents the mean from one experiment. The total numbers of progeny scored in all experiments are indicated in parentheses. (**C**) RNA was purified from dissected gonads and expression levels of mature miRNAs were quantified by qRT-PCR. ***p < 0.001, **p < 0.01, *p < 0.05 (ANOVA with Dunnett’s post-test). N.D., not detected. (**D**) Quantification of the expression of *mir-35–41^prom^::GFP* in embryos whose mothers were exposed to OP50 or CBX102. (**E**) Quantification of the expression of *mir-35–41^prom^::GFP* in WT and *mir-35–41(nDf50)* mutant embryos whose mothers were exposed to CBX102. (**D** and **E**) A.U., arbitrary units. ***p < 0.001 (two-tailed Mann-Whitney test). The numbers of embryos are indicated in parentheses. (**A–E**) Bars represent mean ± SEM.

We next examined zygotic expression of miR-35^fam^ using *wwIs8*[*mir-35–41^prom^::GFP*], a transcriptional reporter driven by the promoter of the *mir-35–41* cluster (Martinez et al., 2008), which is silenced in the maternal germline but active in embryos. The expression of the transcriptional reporter was elevated in embryos whose mothers had experienced CBX102 (Figs. 4D and S7A). Furthermore, the reporter expression was significantly reduced in the *mir-35– 41(nDf50)* mutant background (Fig. 4E). Thus, like the positive feedback-based epigenetic inheritance of chromatin structure (Margueron and Reinberg, 2010), the long-lasting response transmitted from mother to mid-stage embryo may be enabled by an autoregulatory feedback loop of miR-35^fam^ (Fig. 5).

**Figure 5.**
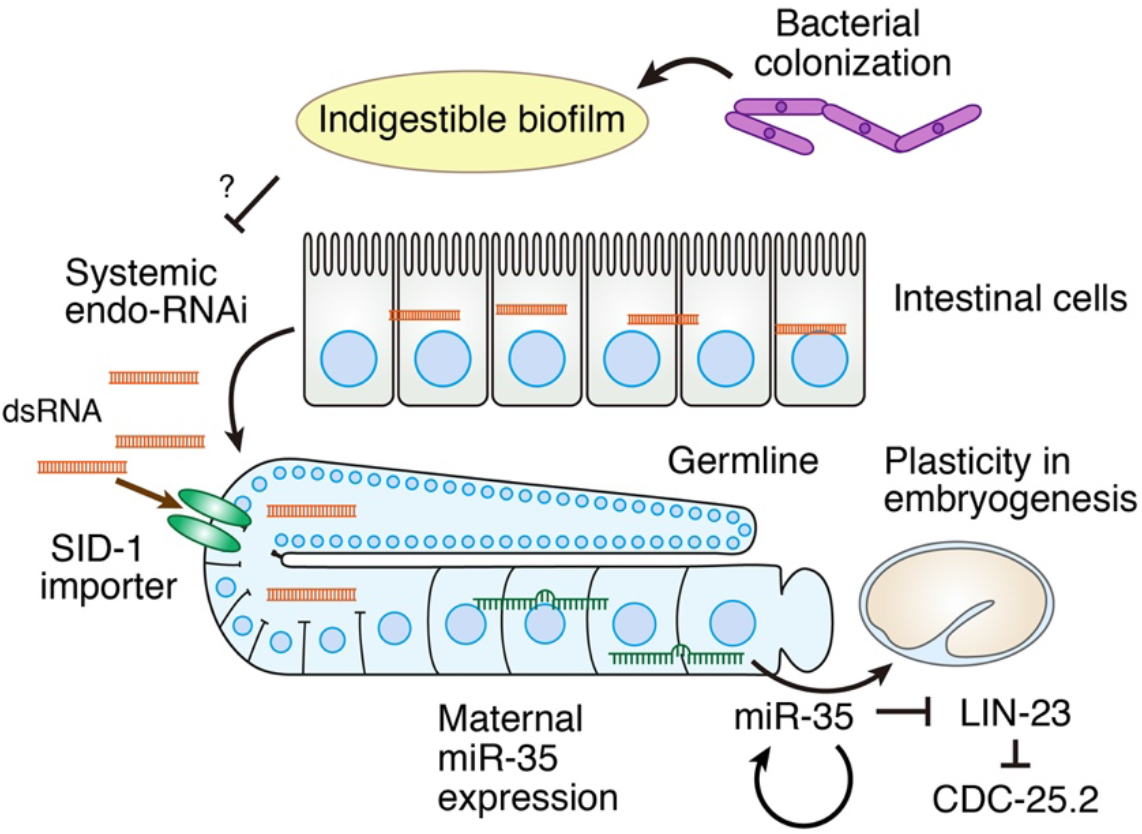
A model for the mechanism of the epigenetic regulation of the alternative embryonic development.

An experimentally verified functional miR-35^fam^-binding site is embedded in the 3’-untranslated region (3’-UTR) of *lin-23* (Liu et al., 2011), which encodes a conserved F-box protein. LIN-23 negatively regulates the intestinal cell number by degrading CDC-25, a family of cell cycle-activating phosphatases, such as CDC-25.2 (Son et al., 2016). To investigate whether miR-35^fam^ regulates extra intestinal cells via the repression of *lin-23*, we used the CRISPR/Cas9 system to generate *lin-23(zb11)*, a short deletion allele in which the miR-35^fam^-binding site in *lin-23* 3’-UTR is disrupted (Fig. S7B). While the *lin-23(zb11)* mutation did not lead to any gross phenotype, it as well as a hypomorphic mutation in *cdc-25.2* suppressed the *nEx1187-induced* extra intestinal cells (Fig. 4B), suggesting that the LIN-23/CDC-25 pathway acts downstream of miR-35^fam^ (Fig. 5). However, the insufficient suppression of extra intestinal cells in *lin-23(zb11)* (Fig. 4B) suggests the potential involvement of other miR-35^fam^ target(s) than *lin-23*. While LIN-23 and CDC-25 are ubiquitously expressed in the embryo with pleiotropic functions, small changes in their activities result in organ-specific abnormalities in intestinal cell divisions (Hebeisen and Roy, 2008; Kostic and Roy, 2002). Our results are consistent with these known phenotypes as well as the notion that miRNAs tune the expression of their targets within a range (Baek et al., 2008; Selbach et al., 2008).

Interactions between animals and microbes in their environment are complex. Some are beneficial, such as maternal gut bacteria in mice providing essential metabolites for embryonic development (Vuong et al., 2020). On the other hand, animals, including humans, are adversely affected by chronic exposure to some indigenous microbes. Given the ability of the microbiome to affect systemic signaling in the host in terms of the nervous, endocrine, and immune systems as well as extracellular vesicles (Sommer and Bäckhed, 2013; Zhang et al., 2018), our study raises the question of whether such exposure could induce adaptive changes in embryos. In *C. elegans*, endo-RNAi is known to repress foreign DNA to protect genome integrity (Billi et al., 2014) and has been proposed as an epigenetic mechanism for adaptation (Grishok, 2013). Our results highlight endo-RNAi as a mechanism to monitor and broadcast pathogen and organ health.

## Materials and Methods

### Strains and culture

Standard methods were used to construct and culture *C. elegans* strains (Brenner, 1974; Dickinson et al., 2013; Frøkjær-Jensen et al., 2012; Mello et al., 1991). Worms were cultured on nematode growth medium (NGM) plates containing 50U/mL nystatin (Sigma Aldrich) at 20–21°C. Bristol N2 was used as the wild type. The strains used in this study are listed in Table S1.

### Embryo mounting and fluorescence microscopy

Preparation and mounting of embryos were performed as previously described (Bao and Murray, 2011) with slight modifications. Briefly, ~20 gravid adult worms were picked to ~20 μl of Boyd buffer (5 mM HEPES [pH 7.2], 3.5 g/L NaCl, 2.4 g/L KCl, 0.4 g/L Na_2_HPO_4_, 0.2 g/L CaCl_2_, 0.2 g/L MgCl_2_, 0.2% glucose) and washed three times with Boyd buffer. Worms were then cut with a 25G needle (BD Precisionglide) and early embryos at the 2- to 30-cell stage were collected with an eyelash. Embryos were transferred with a glass needle and mounted in ~1.5μL of Boyd buffer containing ~50 polystyrene-based microspheres (20 μm in diameter, Polysciences) between a 24 × 50 mm glass slide and an 18 × 18 mm glass coverslip, which was sealed at the edges with melted Vaseline.

Images of live embryos were acquired with an Olympus UPlanSAPO 60× silicone oil objective (NA, 1.30) or an Olympus UPlanSAPO 40× silicone oil objective (NA, 1.25). The microscopes used were (i) a spinning-disk confocal microscope (Quorum Technologies Inc.) consisting of a Zeiss AxioObserver Z1, a Yokogawa CSU-X1 spinning-disk unit, and two Hamamatsu C9100-13 EM-CCD cameras, and (ii) an instant structured illumination microscope (Visitech iSIM) consisting of an Olympus IX73 and a Hamamatsu Flash 4.0v2 sCMOS camera. Total fluorescence of the *mir-35–41::GFP* reporter (Figs. 4D and 4E) was measured in ~30 cell stage embryos using Fiji software (National Institute of Health).

### Exposure of worms to a bacterial lawn

For preculture of bacterial strains, a freshly streaked colony was inoculated and grown at 30°C in LB broth with the exceptions of OG1RF, which was cultured in brain heart infusion (BHI) broth (Difco), and JUb44 and BIGb0170, which were cultured at 25°C. After 24 h, 100 μL of preculture was spread onto a 60 mm NGM plate and then incubated at 20–21°C for 24 h.

NGM plates for culturing parental worms were seeded with OP50 and incubated at 20–21°C for two days. The plates were stored at 4°C for 2 to 20 days until the day of use. Parental worms were cultivated on the OP50-seeded plates until L4 larval to young adult stages. The parental worms were collected and washed five times with M9 buffer (3 g/L KH_2_PO_4_, 6 g/L Na_2_HPO_4_, 5 g/L NaCl, 1 mM MgSO_4_, 0.03% gelatin), and then ~50 worms were transferred to a plate seeded with each indicated bacterial strain. After 24 h of incubation at 20–21°C, their embryos were collected and mounted as described above.

Ivermectin was first dissolved in DMSO at a concentration of 10 μg/mL and added to melted NGM agar to give a final concentration of 1 ng/mL before pouring.

### Quantification of intestinal cell numbers

Numbers of intestinal cells were counted in 1.5- to 3-fold stage embryos using the fluorescence of *stIs10453*[*elt-2^prom^::elt-2::gfp*] or the combination of *rrIs1*[*elt-2^prom^::nls::gfp*] and *ujIs113*[*pie-1^prom^::mCherry::H2B, nhr-2^prom^::mCherry::his-24*]. Because the expression of *rrIs1* was variable and occasionally undetectable in some cells, the additional fluorescence of *ujIs113* was required to accurately count intestinal cells.

Intestinal cell numbers in cross progeny (Fig. S3C) were counted as follows. *him-8(-)* males containing the *stIs10453*[*elt-2^prom^::elt-2::gfp*] marker and *him-8(+)* hermaphrodites lacking any fluorescent marker were raised until mid-L4 stage on OP50-seeded plates. The males and hermaphrodites were separately collected, washed five times with M9 buffer, and transferred to NGM plates on which CBX102 was grown for 18 h. After 6 h of separated preexposure, 50 males and 10 hermaphrodites were picked to a new NGM plate on which CBX102 was grown for 24 h. After 24 h of incubation at 20–21°C, embryos were collected from the hermaphrodites and mounted as described above. Intestinal cells were observed only in the cross progeny, which carried the *stIs10453* marker.

### Cell lineage analyses

For cell lineage analyses, embryos were imaged with 1 μm z-steps across 29 μm and with 75 second intervals. Positions of cell nuclei and timing of cell divisions were determined with StarryNite and AceTree software (Murray et al., 2006). Directions of cell divisions and distance between cells were calculated from the central coordinates of the cell nuclei. 3D reconstruction of cell nuclei positions in embryos (Figs. 2A and 2B) was performed by WormGUIDES software (Santella et al., 2015).

### Brood size and growth assays

L4 and young adult worms (P0) grown on OP50 were collected and washed five times with M9 buffer and then ~50 worms were transferred to a plate seeded with CBX102. After 30 h, 1.5-fold embryos (F1) were directly picked from the CBX102 plate into ~50 μL of M9 buffer dropped on a 24 × 50 mm glass slide, and their ELT-2::GFP fluorescence was observed with the spinningdisk confocal microscope described above. Embryos without extra intestinal cell (E20), embryos with one extra intestinal cell produced by either int7L or int7R divisions (E21), and embryos with two extra intestinal cells produced by both int7L and int7R divisions (E22) were isolated. The F1 embryos were individually plated onto NGM plates seeded with OP50 or onto NGM plates seeded with a mixture of stationary phase cultures of CBX102 and OP50 mixed in a volumetric ratio of 1:9 (10% CBX102 plates).

For brood size assays, the isolated F1 embryos were grown until young adults and all of the F2 hatchlings were killed and counted every 24 h until the F1 worms no longer produced progeny. For growth assays on CBX102 (Fig. S6C), the isolated F1 embryos were allowed to develop at 20°C for 80 h on 10% CBX102 plates and their developmental stages were determined.

For growth assays on the *E. coli* and *B. subtilis* strains (Fig. S3E), early embryos were prepared by hypochlorite treatment of gravid hermaphrodites raised on OP50 and were transferred onto a new plate seeded with an indicated bacterial strain. The embryos were allowed to develop at 20°C for 65 h and their developmental stages were determined.

### Isolation of gonadal miRNAs and RT-qPCR

Young adults were transferred into Egg buffer (25 mM HEPES [pH 7.2], 118 mM NaCl, 48 mM KCl, 2 mM CaCl_2_, 2 mM MgCl_2_) and washed five times with Egg buffer immediately before dissection. The gonads were isolated from one hundred worms by cutting the worms behind the pharyngeal bulb and in front of the spermatheca with a 25G needle. The isolated gonads were placed into 300 μL TRIzol on ice. Sixty microliters of chloroform were added to it, and after centrifugation, the aqueous phase was transferred into a new tube. After adding 1.5 volumes of ethanol, the samples were loaded into a miRNeasy column (Qiagen) and purified according to the supplier’s instructions. The purified samples were reverse-transcribed using the miSctipt reverse transcriptase and HiSpec buffer (Qiagen). For quantitative PCR, the Quantitect SYBR Green PCR kit (Qiagen) was used on the QuantStudio 6 Flex real-time PCR system (Applied Biosystems). Primers specific for miR-2-5p, miR-47-3p, miR-35-3p, mir-40-3p, and miR-42-3p were obtained from the miScript Primer Assays kit (Qiagen). miR-47-3p was used as the endogenous control, and the results were analyzed using the comparative Ct method.

### Germline transformation

Expression constructs were injected at 10–50 ng/μL. pG-*unc-122^prom^::mCherry* and pCFJ90 (*myo-2^prom^::mCherry*, a gift from Erik Jorgensen, Addgene plasmid #19327) were utilized as injection markers. In each case, the total concentration of injected DNA was 100 ng/μL.

The *zb11* allele of *lin-23* was generated using the CRISPR-Cas9 system (Dickinson et al., 2013). The sgRNA expression construct containing the target sequence 5’-GGTTTGGTTGATTTCTGCAC-3’ was obtained using a PCR-fusion technique and the PCR product was injected at 15 ng/μL together with 50 ng/μL pDD162 (*eft-3^prom^::Cas9*, a gift from Bob Goldstein, Addgene plasmid #47549). After injection, F1 animals were screened for a deletion by single-worm PCR using the primers 5’-TATGGATCACCTGGGCGGAG-3’ and 5’-GAGGAAAAGTTGGGAAGGGGA-3’. The resulting *lin-23(zb11)* mutant line was backcrossed twice to the original strain before use.

The *pie-1^prom^::sid-1(cDNA)::pie-1 3’UTR* transgene in the *pCFJ150-pie-1^prom^-sid-1(cDNA)-pie-1 3’UTR* plasmid (see below) was integrated as a single copy into the *oxTi185* site on chromosome I using the standard MosSCI technique (Frøkjær-Jensen et al., 2012).

### Transgene constructions

The construction of pDEST-*Cas9*, pENTR-*ges-1^prom^*, pENTR-*myo-3^prom^*, pENTR-*rimb-1^prom^*, and pG-*unc-122^prom^::mCherry* was described previously (Ohno et al., 2014, 2017a, 2017b). pG-*ges-1^prom^-Cas9*, pG-*rimb-1^prom^-Cas9*, and pG-*myo-3^prom^-Cas9* were made by LR recombination reactions between the pENTR plasmids and pDEST-*Cas9*. For pCFJ150-*pie-1^prom^-sid-1(cDNA)-pie-1 3’UTR*, the *sid-1* cDNA was amplified from the genomic DNA of the XE1375 strain (Firnhaber and Hammarlund, 2013) with the primers 5’-gtgtAGGCCTaaaaATGATTCGTGTTTATTTGATAATTTTAATGCA-3’ and 5’-gtgtGCTAGCCTAGAAAATGTTAATCGAAGTTTTGCGT-3’ and inserted into the *Nae*I-*Nhe*I sites of pCFJ150-*mCherry(dpiRNA)::ANI-1(AHPH)* (a gift from Heng-Chi Lee, Addgene plasmid #107939). Three expression constructs of P*U6::rrf-3 sgRNAs* were amplified by PCR from pDD162 using the primers 5’-GTATTGTGTTCGTTGAGTGACC-3’, 5’-TGGCTTAACTATGCGGCATC-3’, 5’-gGAAATGTTGATTGACTCTGgttttagagctagaaatagc-3’, 5’-CAGAGTCAATCAACATTTCcaagacatctcgcaataggag-3’, 5’-gAGAAAACGAAGCACAGAGGgttttagagctagaaatagc-3’, 5’-CCTCTGTGCTTCGTTTTCTcaagacatctcgcaataggag-3’, 5’-gCTCAAATCTCGCATACGAGgttttagagctagaaatagc-3’, and 5’-CTCGTATGCGAGATTTGAGcaagacatctcgcaataggag-3’ (target sequences were GGAAATGTTGATTGACTCTG, GAGAAAACGAAGCACAGAGG, and GCTCAAATCTCGCATACGAG), and were co-introduced. Further details of the expression constructs will be provided upon request.

### Statistical analyses

Statistical analyses were performed with a statistic package (Prism v.5, GraphPad software). Error bars indicate SEM. All experiments were repeated three to seven times with similar results.

## Acknowledgments

We thank the *Caenorhabditis* Genetics Center (funded by the NIH Office of Research Infrastructure Programs P40 OD010440), NBRP (NIG, Japan)—*B. subtilis*, NBRP (Tokyo Women’s Medical University, Japan)—*C. elegans*, the *Bacillus* Genetic Stock Center, Dr. Jeremy Nance, and Dr. Danielle A. Garsin for strains.

## Author contributions

H.O. and Z.B. conceived and designed the study and wrote the manuscript; H.O. performed the experiments and did the analyses.

## Data and materials availability

The raw data will be deposited on Figshare. All strains and plasmids generated in this study will be available from the authors upon reasonable request.

---

**Figure S1.**
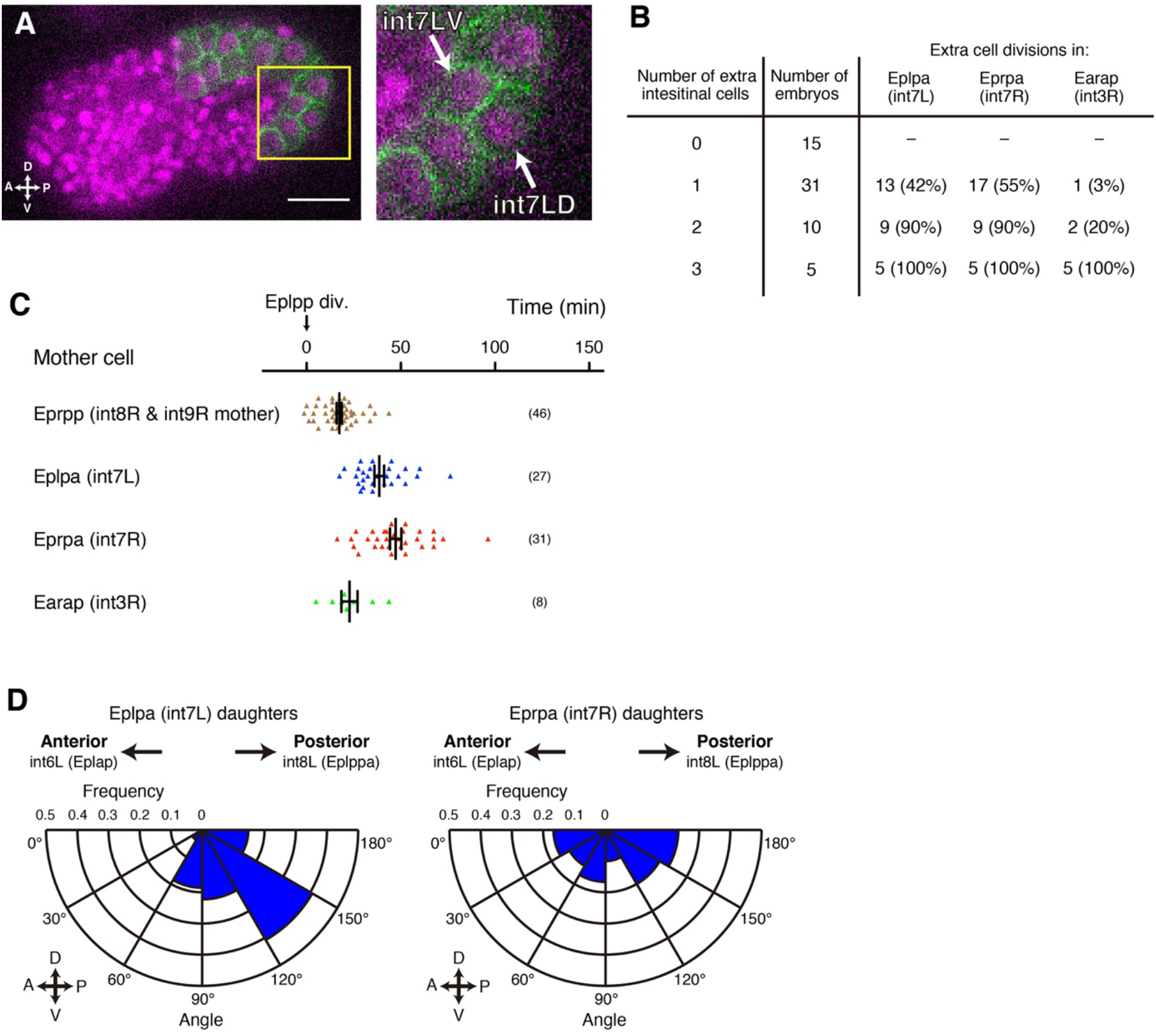
Extra cell divisions induced by CBX102 occur at specific cells. (**A**) Representative embryo with one extra intestinal cell, whose mother was exposed to CBX102. Green, intestinal cell membranes labeled with *end-1^prom^::gfp::caax*; magenta, all cell nuclei labeled with mCherry::Histone. A magnified view of the boxed region (yellow) is shown in the right panel. The two nuclei (white arrows) produced by the extra division of int7L were surrounded by the cell membranes. Scale bar, 10 μm. (**B**) Numbers of extra intestinal cells and numbers of embryos that underwent extra divisions in Eplpa (int7L), Eprpa (int7R), and Earap (int3R), among 61 embryos whose mothers were exposed to CBX102. (**C**) Timing of extra cell divisions of Eplpa (int7L), Eprpa (int7R), and Earap (int3R) in 46 embryos whose mothers were exposed to CBX102. The division of Eplpp, the mother of int8L and int9L, was set to time zero. The timing of the division of Eprpp (the mother of int8R and int9R), which occurred in all of the embryos, is also shown. The numbers of cell divisions observed in the 46 embryos are indicated in parentheses. (**D**) Frequency distribution of the axial direction of the extra cell divisions of Eplpa (int7L, left) and Eprpa (int7R, right). The line drawn between the centers of int6L and int8L cell nuclei is regarded as the antero-posterior (AP) axis. n = 27 divisions for Eplpa and 30 divisions for Eprpa.

**Figure S2.**
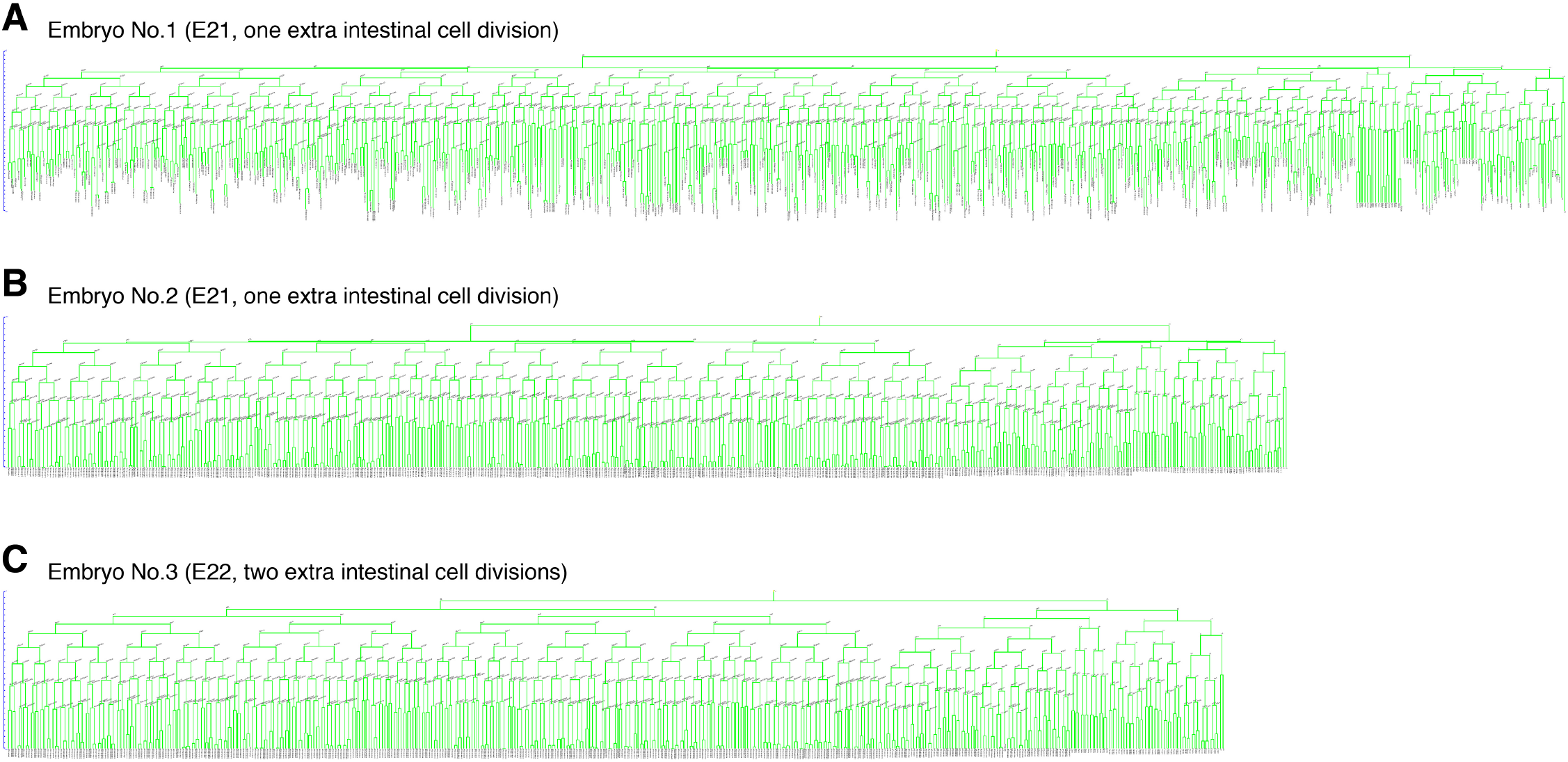
Gut microbes specifically affect the intestinal cell lineage. (**A**) Cell lineage tree, determined by extensive tracking of cell nuclei, of an embryo with one extra intestinal cell. A horizontal line in the lineage tree indicates a cell division. No abnormal cell division nor abnormal cell death event was detected in any lineage other than the E lineage until 555 cells (563 cell divisions and nine cell deaths). Normal occurrence of additional 103 cell divisions and 78 cell deaths was confirmed by eye, covering 99.4% (666/670) of the cell divisions and 77% (87/113) of the cell deaths that occur during embryogenesis. (**B**) Cell lineage tree of another embryo with one extra intestinal cell up to 533 cells. No abnormal cell behavior was observed except for an extra division of Eprpa (int7R). (**C**) Cell lineage tree of an embryo with two extra intestinal cells up to 506 cells. No abnormal cell behavior was observed except for extra divisions of Eplpa (int7L) and Eprpa (int7R).

**Figure S3.**
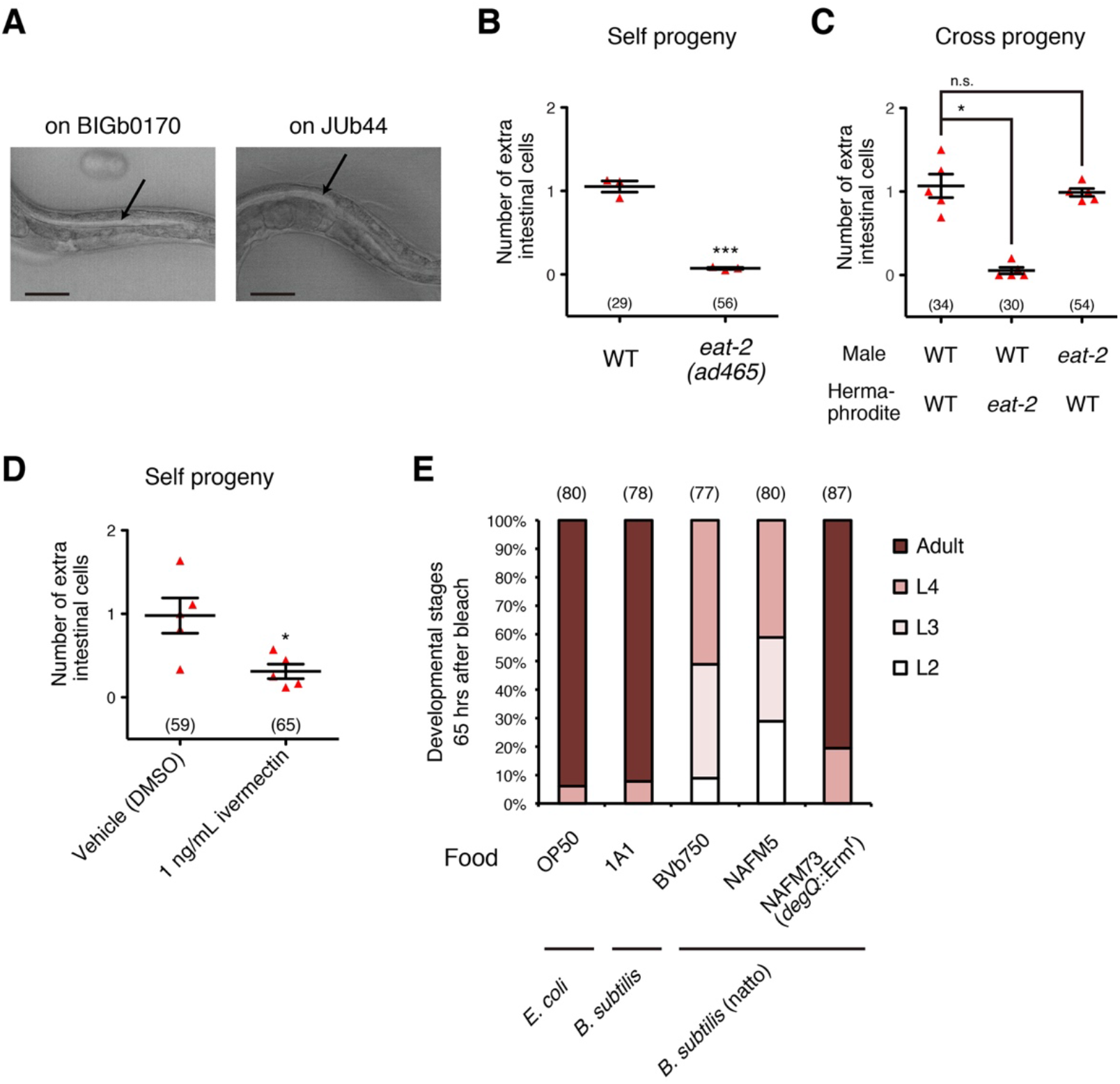
Maldigestion of maternal gut microbes may trigger the developmental plasticity. (**A**) Representative photomicrographs of WT adults which were incubated on BIGb0170 or JUb44 for 24 h. Arrows, intestinal lumen. Scale bars, 50 μm. (**B**) Numbers of extra intestinal cells in self-progeny of WT and feeding-defective *eat-2(ad465)* mutants. The mothers were exposed to CBX102. ***p < 0.001 (two-tailed *t*-test). (**C**) Numbers of extra intestinal cells in cross progeny of WT and *eat-2(ad465)* whose parents were exposed to CBX102. *p < 0.05 (Kruskal-Wallis and Dunn’s post-test). n.s., not significant. (**D**) Numbers of extra intestinal cells in progeny whose mothers were exposed to CBX102 together with the anthelmintic drug ivermectin, which inhibits pharyngeal pumping. 0.01% DMSO was used as vehicle control. *p < 0.05 (two-tailed *t*-test). (**B–D**) Bars represent mean ± SEM of at least three independent experiments. Each dot represents the mean from one experiment. The total numbers of progeny scored in all experiments are indicated in parentheses. (**E**) Fractions of developmental stages of wild-type worms cultured with *E. coli* OP50, *B. subtilis* 1A1, *B. subtilis (natto)* BVb750 and NAFM5, and γPGA-nonproducing *B. subtilis* (natto) NAFM73. Bleached embryos were transferred to a new seeded plate and allowed to develop at 20°C for 65 h. The numbers of worms tested are indicated in parentheses.

**Figure S4.**
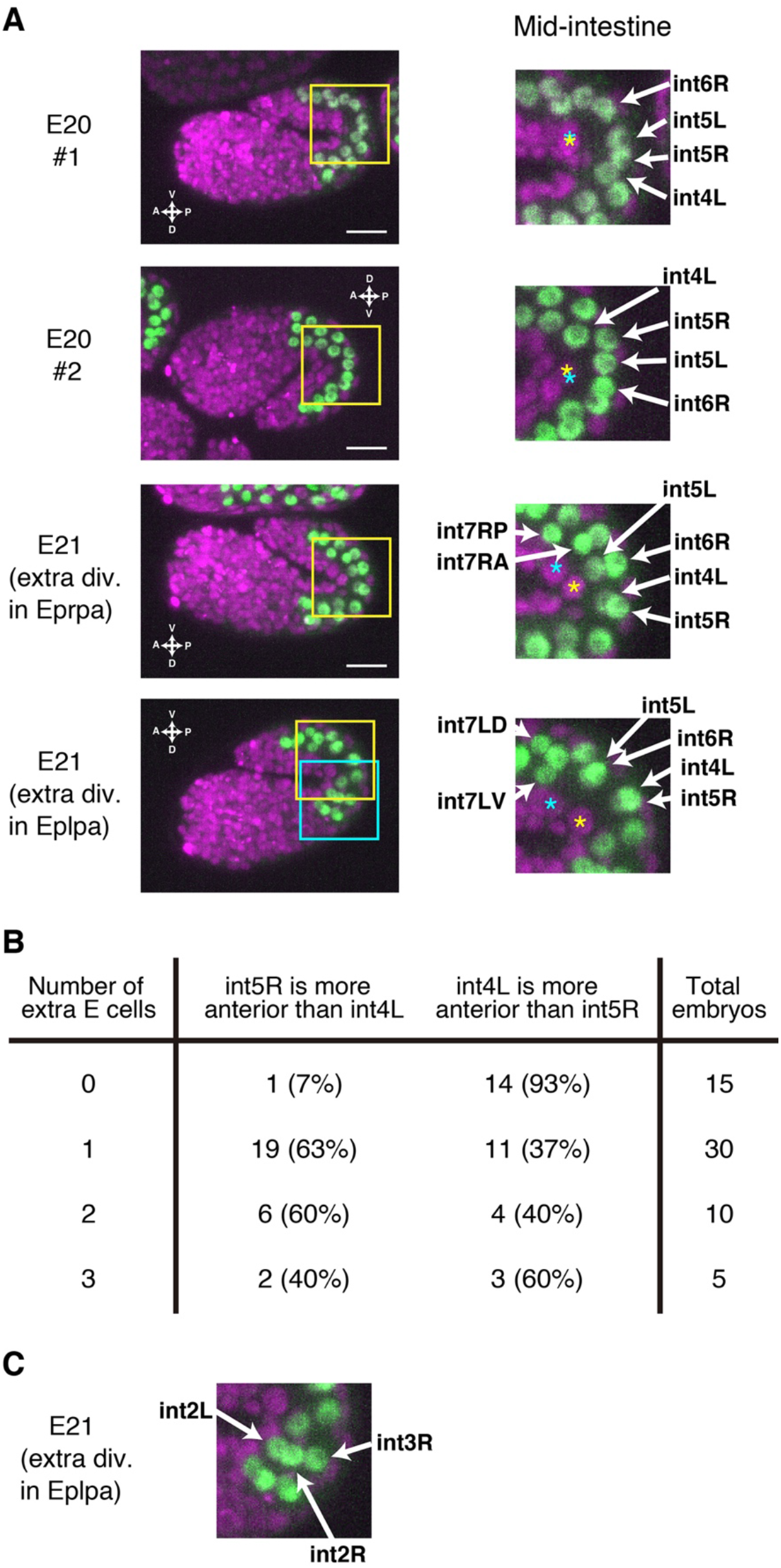
Extra intestinal cells coincide with left-right asymmetric cell pairing. (**A**) Two representative embryos without extra intestinal cell (E20) and two representative embryos with one extra intestinal cell (E21), whose mothers were exposed to CBX102. Green, intestinal nuclei labeled with *elt-2::gfp;* magenta, all nuclei labeled with mCherry::Histone. Magnified images of the boxed regions (yellow) are shown in the right panels. Yellow asterisks, Z2; cyan asterisks, Z3. Scale bars, 10 μm. (**B**) Fractions of embryos in which int5R was more anterior than int4L (left) and ones in which int4L was more anterior than int5R (right). (**C**) A magnified view of the boxed region (cyan) in one of the embryos with one extra intestinal cell shown in (A, bottom).

**Figure S5.**
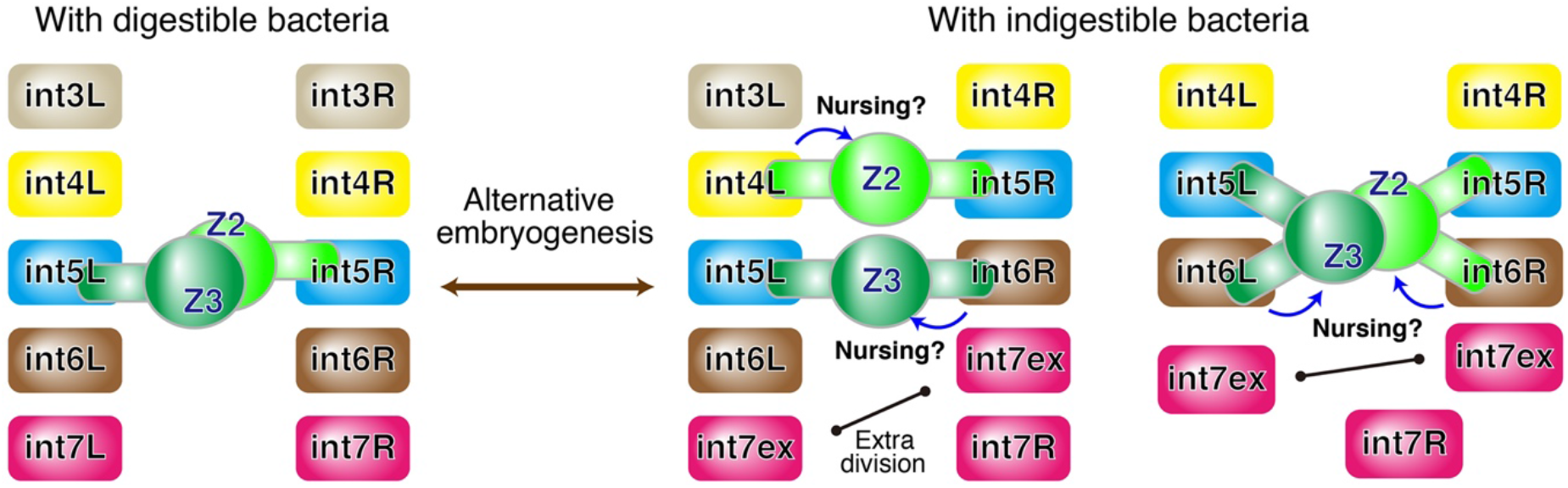
Simplified schematic of the positional changes of the mid-intestinal cells and the increased PGC-intestine association.

**Figure S6.**
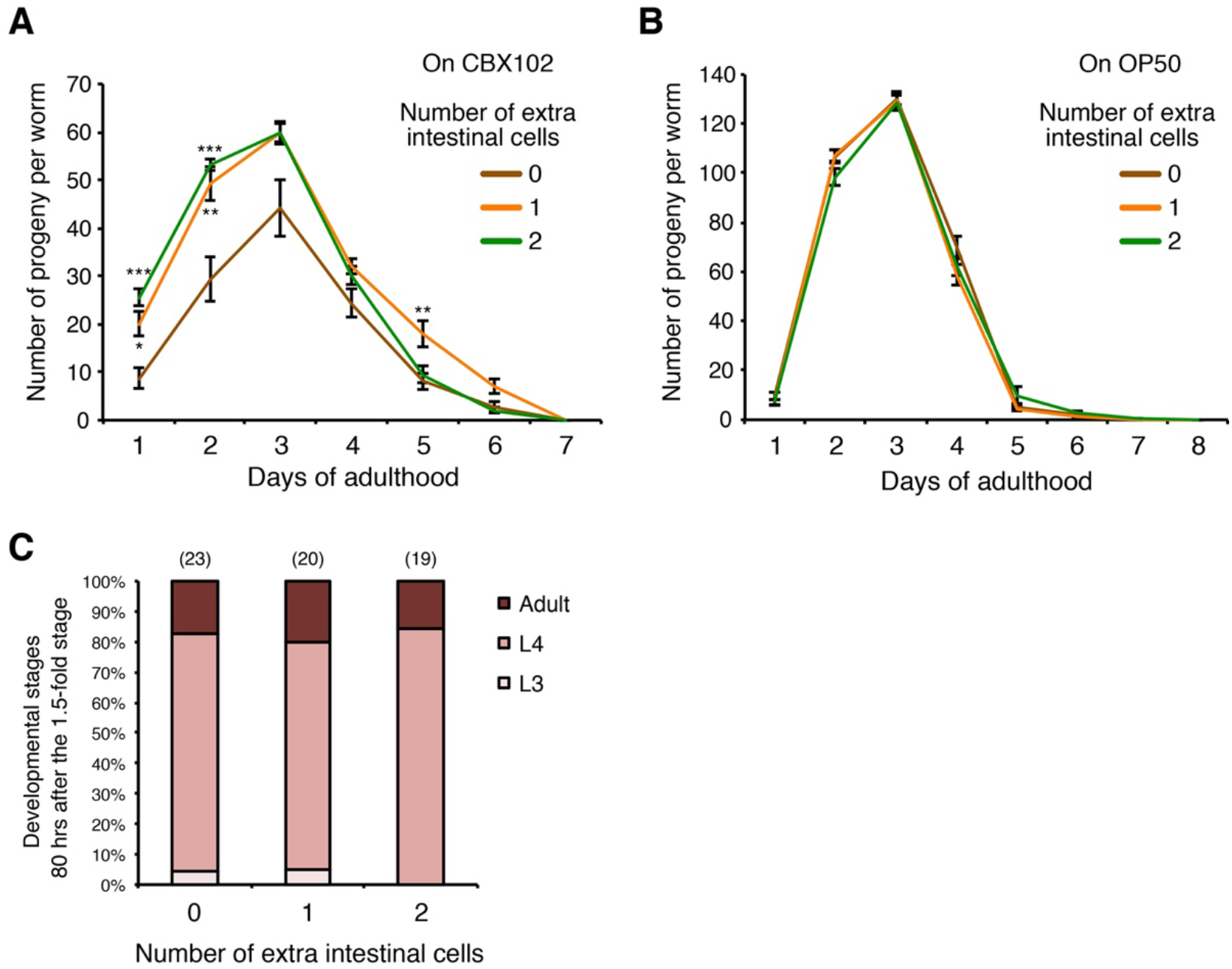
Extra intestinal cells may improve fitness in the presence of harmful microbes. (**A** and **B**) Daily progeny production of the worms shown in Figs. 2G and 2H. Bars represent mean ± SEM. ***p < 0.001, **p < 0.01, *p < 0.05 (Kruskal-Wallis and Dunn’s post-test). (**C**) Fractions of developmental stages of worms cultured with CBX102. Intestinal cell numbers were determined in 1.5-fold stage embryos whose mothers had been exposed to CBX102, and the embryos were individually plated and allowed to develop at 20°C for 80 h. The numbers of worms tested are indicated in parentheses.

**Figure S7.**
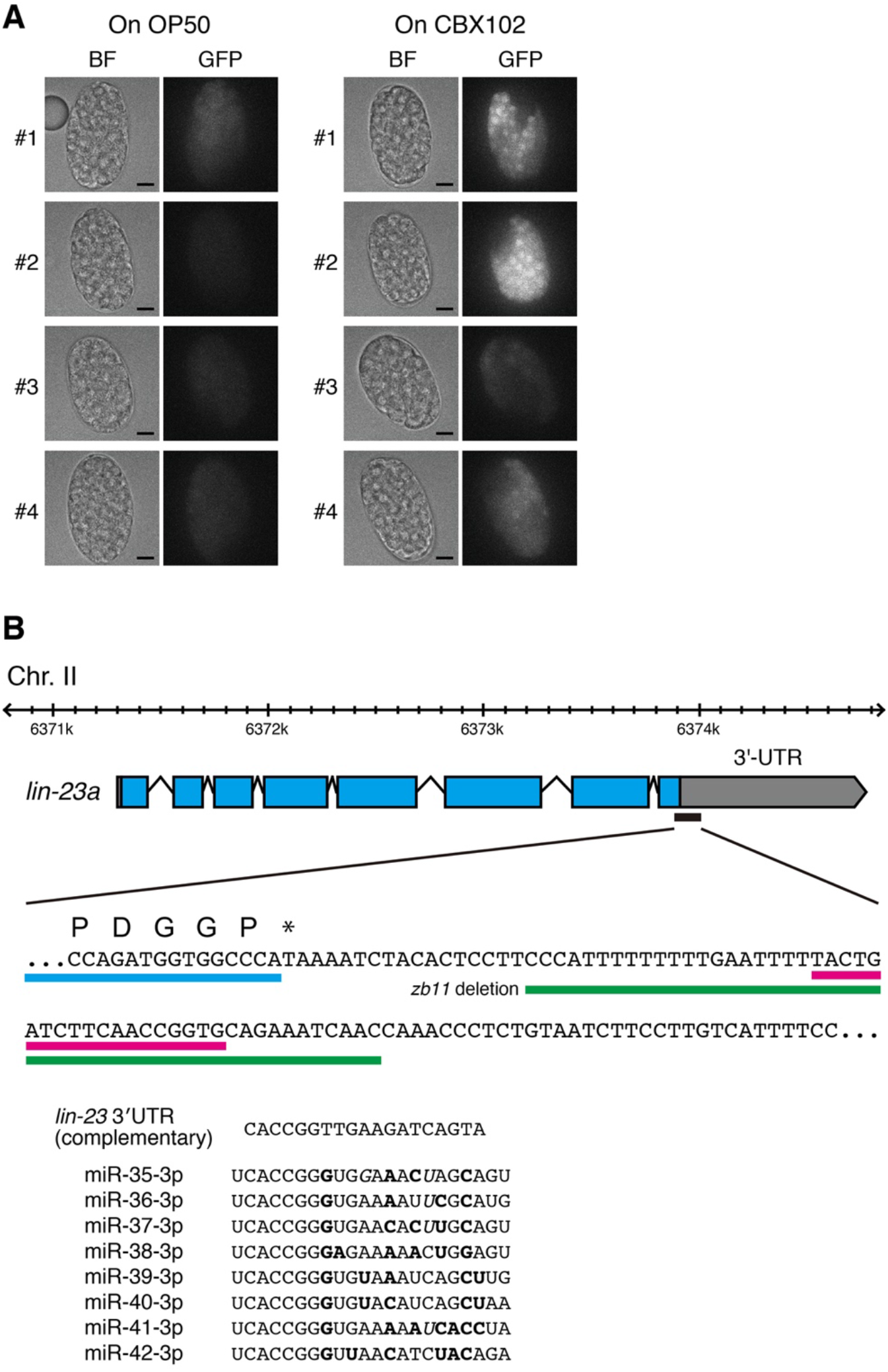
The promoter activity of the *mir-35–41* cluster is enhanced on CBX102. (**A**) Representative embryos of the strain expressing *mir-35–41^prom^::GFP*. Their mothers were exposed to OP50 or CBX102. Scale bars, 10 μm. (**B**) Genomic organization of the *lin-23* locus (top) and sequence alignment of the miR-35^fam^-binding site in *lin-23* 3’-UTR and the mir-35^fam^ miRNAs (bottom). Blue, *lin-23* exons; red, miR-35^fam^-binding site; green, lesion in the *zb11* allele; bold, mismatches; italic, G:U wobbles.

**Table S1.**
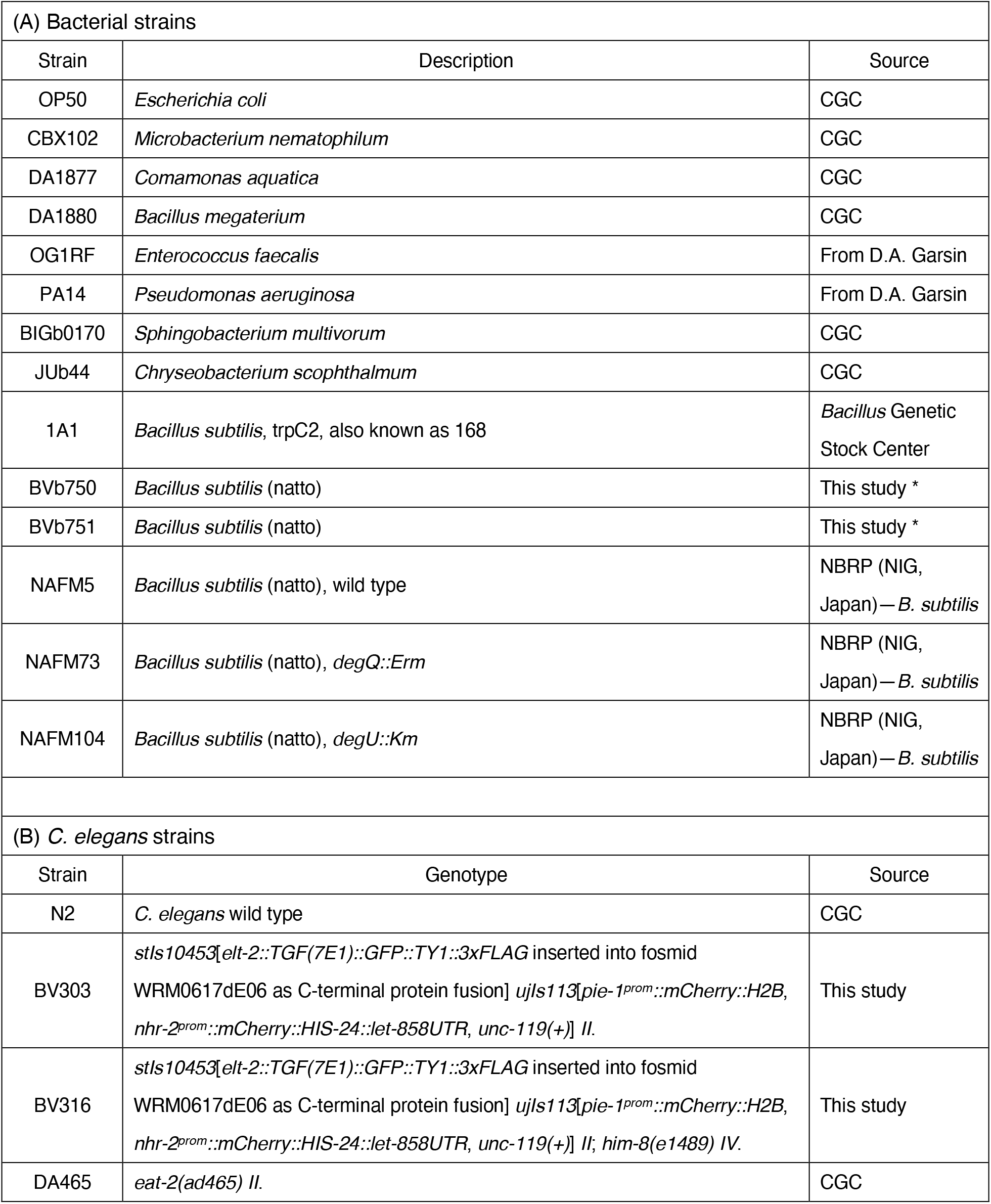
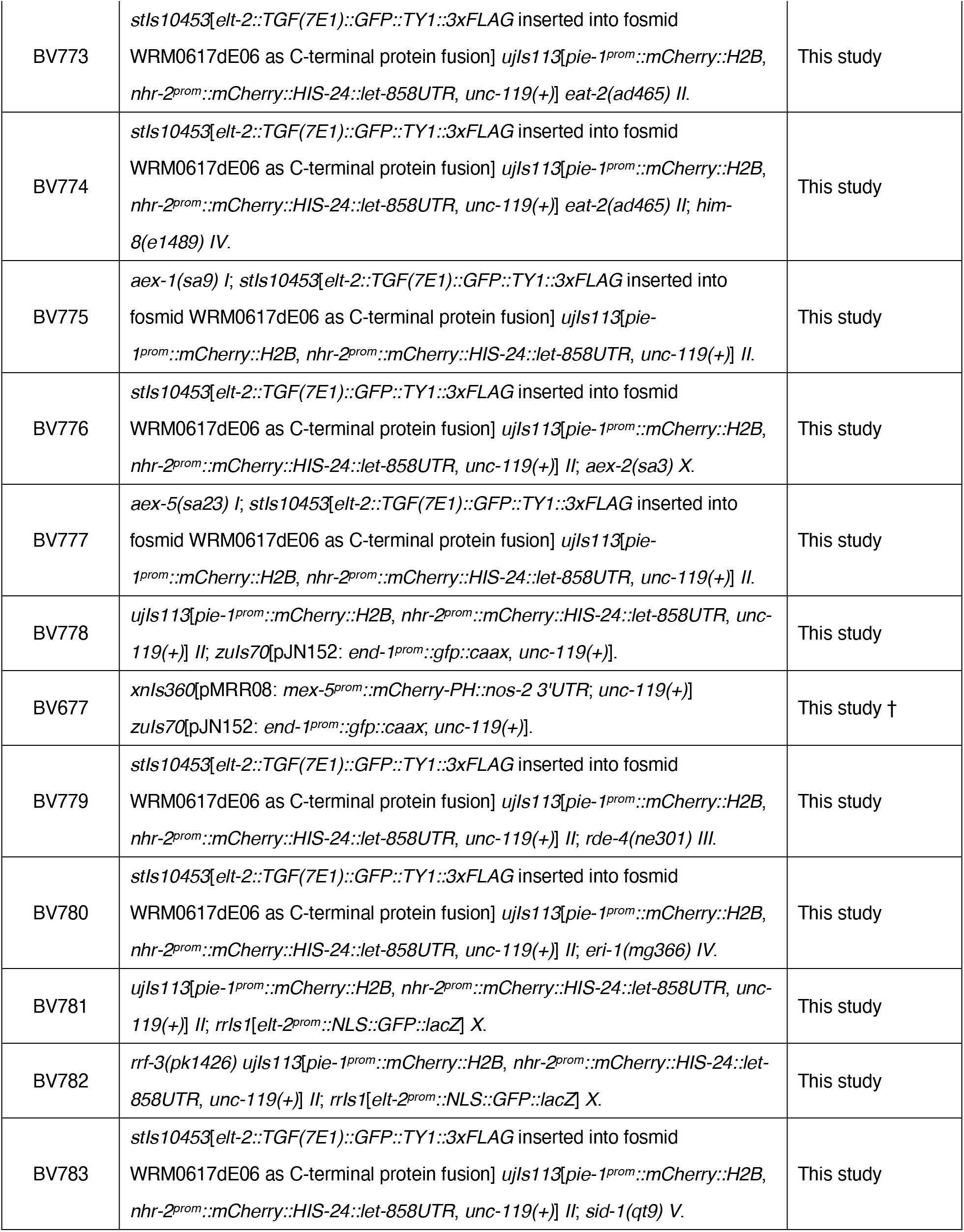
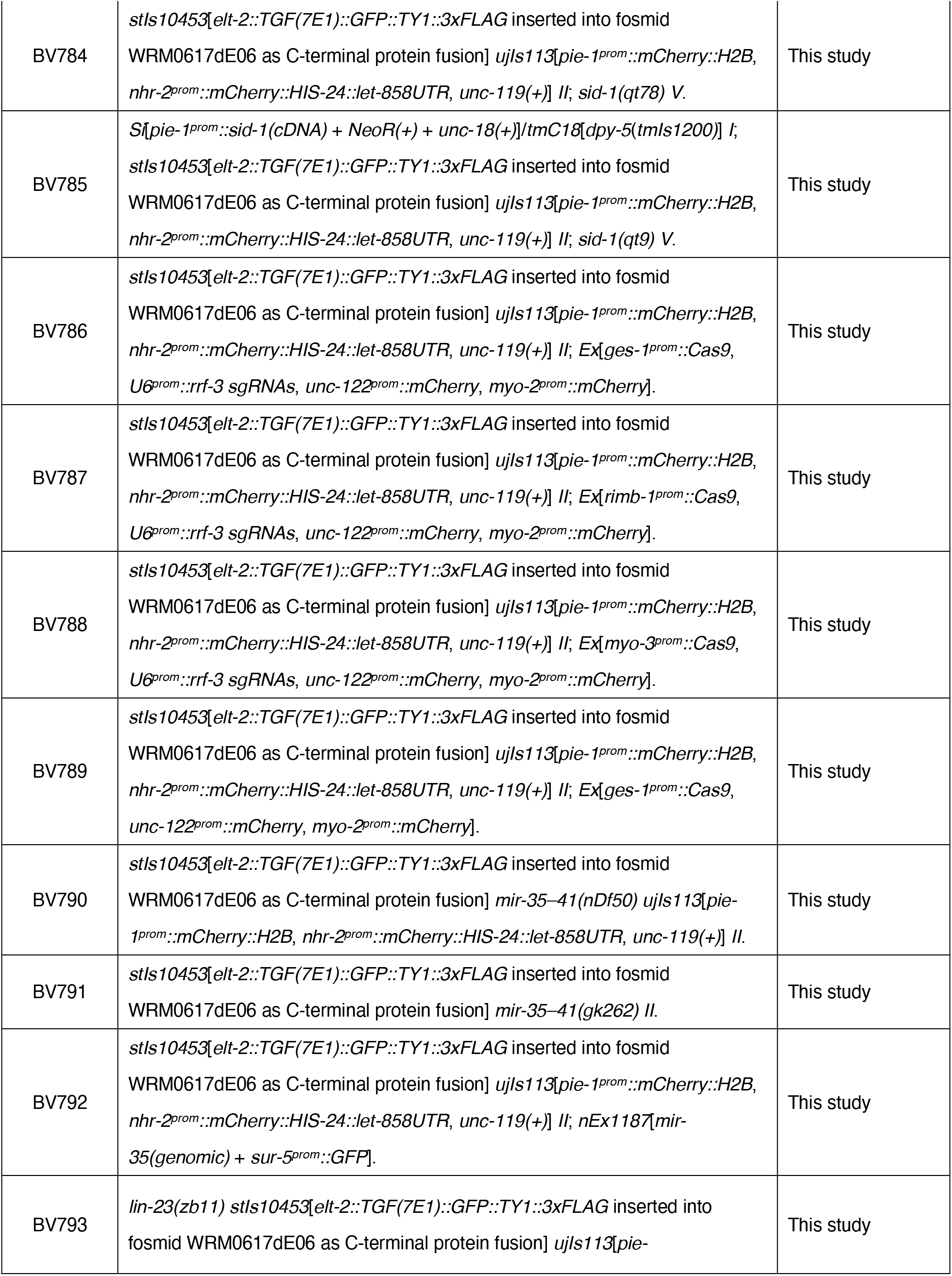
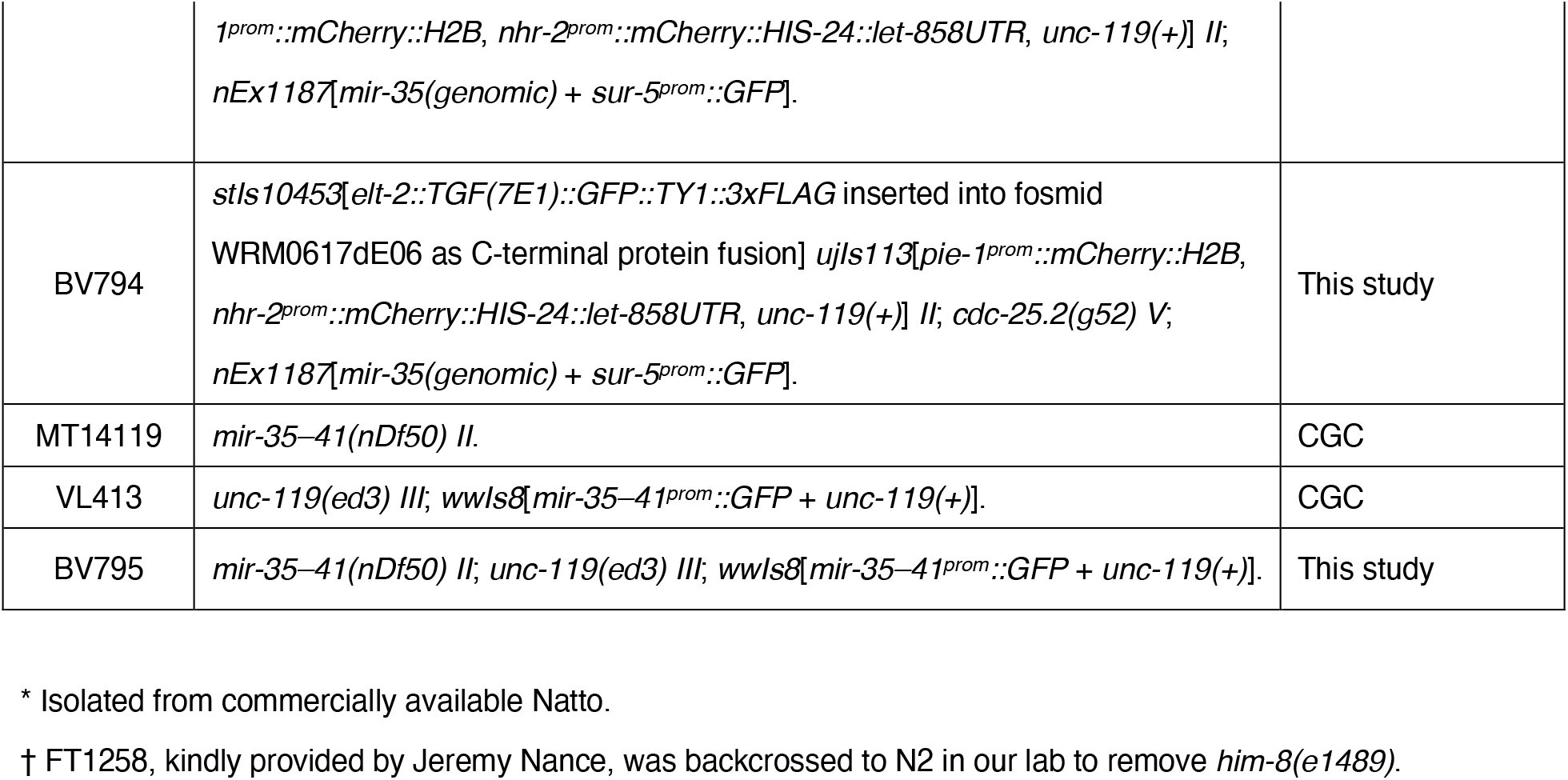
Strains used in this study.

